# Broadband amplification of organ of Corti vibrations in the mouse cochlear apex

**DOI:** 10.64898/2026.06.02.729685

**Authors:** James B. Dewey

**Affiliations:** Caruso Department of Otolaryngology – Head & Neck Surgery, Keck School of Medicine, University of Southern California, Los Angeles, CA 90033, USA

**Keywords:** cochlear mechanics, outer hair cell motility, optical coherence tomography

## Abstract

Mammalian hearing depends on the active amplification of sound-evoked waves as they travel along the basilar membrane within the cochlea. This amplification is mediated by the outer hair cells (OHCs), which generate force to enhance the vibrations of the surrounding structures. While OHCs at a given location only amplify basilar membrane motion for a narrow frequency range, recent measurements show that the amplification of motions deeper within the organ of Corti is much more broadband. However, the extent to which this broadband amplification influences the motions that are most relevant to inner hair cell stimulation – i.e., at the organ’s apical surface – remains uncertain. Here, optical coherence tomography was used to demonstrate that OHCs nonlinearly amplify the motions near the top of the organ of Corti, including at the reticular lamina and tectorial membrane, over a wide frequency range in the mouse cochlear apex. Responses at all frequencies were physiologically vulnerable and grew compressively with stimulus level. Low-frequency responses also exhibited non-monotonic features that were due to interference between amplified motion and the underlying traveling wave. The data suggest that broadband amplification of motions at the top of the organ of Corti likely explains certain phenomena observed in auditory nerve responses.

## INTRODUCTION

The sensitivity, dynamic range, and frequency selectivity of the mammalian auditory system all depend on an active amplification process mediated by the cochlear outer hair cells (OHCs)^1^. OHCs first detect sound-evoked waves traveling along the underlying basilar membrane (BM) via deflection of their stereocilia, which gates mechanotransduction channels and modulates ionic current flow^2^. The resulting changes in membrane potential then drive the motor activity of prestin, a voltage-sensitive protein densely expressed in the OHC lateral wall, causing the cells to change length and generate force^3–6^. These electromotile forces act locally to influence the motion of the surrounding structures and, through mechanisms still being investigated, dramatically amplify BM-coupled traveling waves^7–12^. Determining how local force transmission and the spatially distributed process of traveling-wave amplification interact is critical to understanding how OHCs enhance and tune the stimulus to the inner hair cells (IHCs), the primary afferent sensory receptors communicating with the auditory nerve^13^.

Due to the BM’s mechanical properties, each location along the cochlear spiral is tuned to a particular characteristic frequency (CF), such that low-frequency waves peak at the cochlear apex and high-frequency waves peak more basally^14^. For high-CF locations, it is well established that OHC-generated forces amplify traveling-wave motion over a narrow region near the wave’s peak, such that BM motion at a given location is only amplified at frequencies within an octave or so of the CF^1,15–18^. However, recent advances in optical techniques such as optical coherence tomography (OCT) have allowed measurement of vibrations deeper within the organ of Corti and revealed that the motions near the OHC region are often amplified over a much wider frequency range^18–24^. OHCs therefore appear to be stimulated and generating force in a broadband manner^25^, amplifying their own motion but not the motion of the BM at frequencies far below the CF.

At these low frequencies, the motion near the center of the organ of Corti has most consistently exhibited the physiological vulnerability and nonlinear, compressive growth with stimulus level that indicate the influence of OHC-mediated amplification^21,24,26,27^. However, these features have been more variable in measurements near the top of the organ of Corti, at the reticular lamina (RL) or the overlying tectorial membrane (TM)^19,20,22,26,27^. Although previous OCT measurements of RL and TM motion in the mouse cochlear apex have shown below-CF nonlinearity^18,28,20,29^, concerns regarding imaging resolution^26^ and inconsistencies in the motion phases across reports^18,20,24^ somewhat cloud interpretation of these data. Detailed spatial analysis of the motions in the mouse apex using an improved OCT system^24^ has also indicated that locations previously identified as being near the RL^18^ were likely closer to the lateral supporting cells or deeper in the OHC region. The extent to which below-CF OHC force generation influences the motions that are most relevant to stimulation of the IHCs therefore remains uncertain^13,30^.

Here, insights gained from the previous spatial analysis of motions in the mouse cochlear apex^24^ were used to more confidently identify anatomical locations and measure their sound-evoked displacements. Responses were examined over a wide range of stimulus frequencies and levels to unequivocally demonstrate that OHCs provide broadband amplification of the motions near the top of the organ of Corti, including at the RL and TM. The data provide a comprehensive view of the relation between active OHC-driven motions and the underlying traveling wave and suggest that certain phenomena observed in auditory nerve responses may be explained by broadband amplification of RL and TM motion.

## RESULTS

### Broadband nonlinear amplification influences the motion of all locations beyond the BM

OCT was used to image the mouse cochlear apex *in vivo* and measure sound-evoked displacements from the BM, the bottom of the OHC region near the OHC-Deiters’ cell (DC) junction, the top of the OHC region near the RL, the TM, and a point at the top of the organ of Corti located near the border between the OHCs and the lateral supporting cells, which is referred to here as the lateral cell (LC) region. **Figure 1** shows the locations of these measurement points in the OCT images (**Fig. 1a-c**) as well as their displacement in response to tones varied in frequency and sound pressure level (SPL) for an individual mouse (top panels of **Fig. 1d-h**).

**Figure 1.**
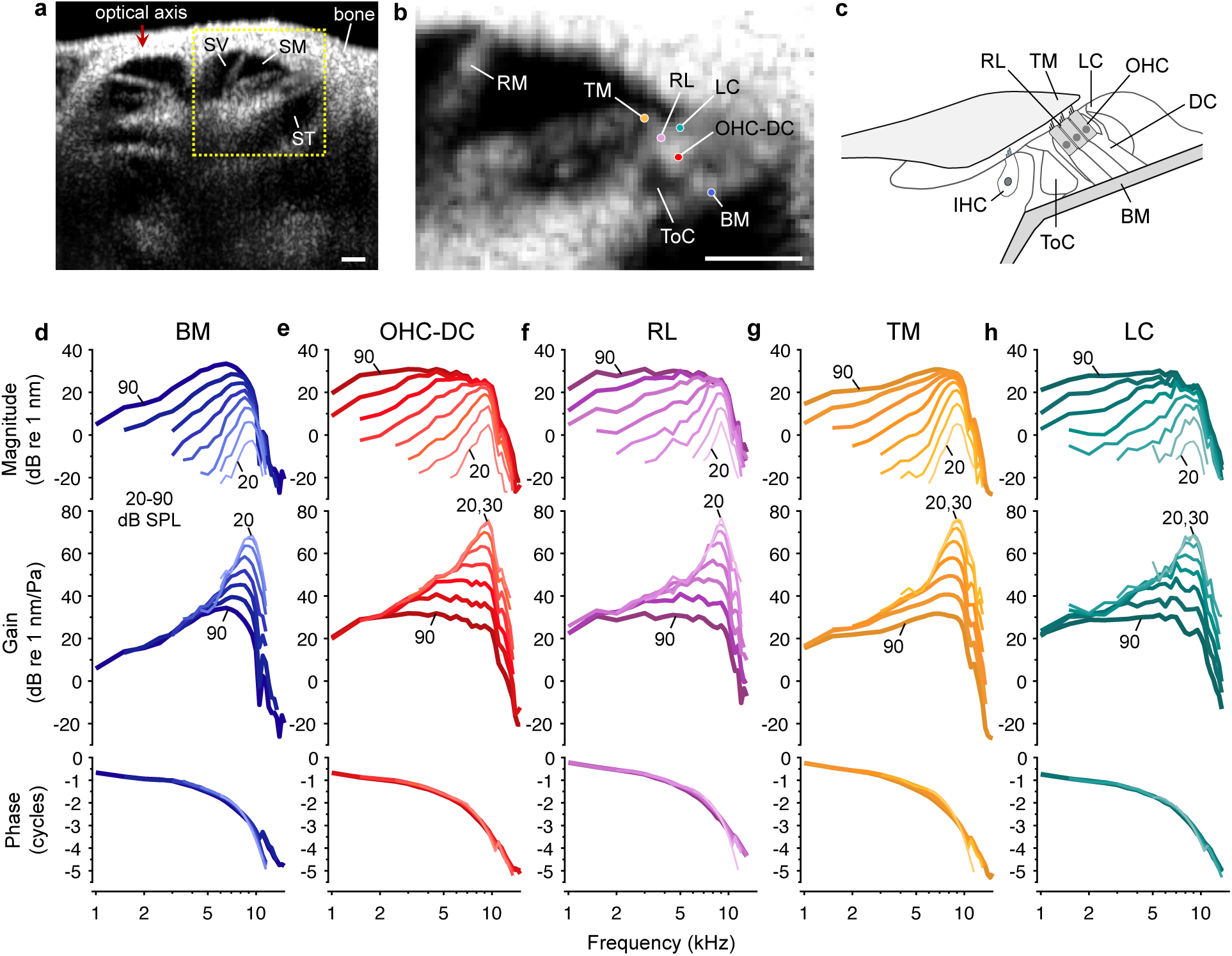
Nonlinearity in sound-evoked displacements extends to lower frequencies for all locations beyond the BM. (**a**) Cross-sectional OCT image of the mouse cochlea. The apical region where displacement measurements were obtained is highlighted in yellow. The three fluid-filled scalae are also indicated (ST = scala tympani; SM = scala media; SV = scala vestibuli). Scale bar = 100 μm. (**b**) Close-up image of the apical region illustrating the locations of specific measurement points on the BM, OHC-DC junction, RL, TM, and LC region. Reissner’s membrane (RM) and the tunnel of Corti (ToC) are also labeled. (**c**) Schematic of the organ of Corti and surrounding structures. (**d**-**h**) Displacement responses to single tones measured as a function of stimulus frequency and level for the BM, OHC-DC junction, RL, TM, and LC region in an individual mouse. Displacement magnitudes (top panels) are shown along with the displacements normalized to the evoking stimulus pressure (i.e., the displacement “gain”; middle panels) and the phases referenced to the stimulus phase measured in the ear canal (bottom panels). Responses to higher-level stimuli are shown with thicker and darker lines, and responses to the lowest and highest levels are indicated numerically. While BM response gains remain nearly constant across stimulus level below ∼4 kHz, all other measurement locations exhibit nonlinear gain down to 2 kHz or lower.

At low stimulus levels, displacements of all locations were sharply tuned to a CF of ∼9 kHz. As the stimulus level was increased, responses grew compressively at frequencies near the CF but grew more linearly at lower frequencies. Due to this frequency-dependent compression, displacements for high-level stimuli became more broadly tuned and tended to peak at lower frequencies. This was particularly true for locations beyond the BM, which moved more at low frequencies and exhibited compressive growth over a broader frequency range. Nonlinear compression in the motions both at and below CF is thought to result from the saturation of OHC mechanotransduction currents – and thus the drive to electromotility – with increasing stimulus level^2^.

To better visualize the nonlinearity, displacements were divided by the evoking stimulus pressure (in Pascals) to calculate the “gain” of the responses (middle panels in **Fig. 1d-h**). Near the CF, the gain for all locations was highest at low stimulus levels and decreased at higher stimulus levels, reflecting the compressive growth in the displacements. However, while BM responses became essentially linear at frequencies more than an octave below the CF, as revealed by the overlapping gain curves, the responses of all other locations showed nonlinearity down to at least 2 kHz, indicating the influence of OHC-mediated amplification.

Displacement phases for all locations exhibited increasing lags with frequency that were consistent with traveling-wave propagation (bottom panels of **Fig. 1d-h**). Systematic phase shifts with level were evident, most notably for the TM at frequencies between the CF and an octave lower, though these are relatively subtle at the scale shown. There were also marked differences in the phases of the motions of certain locations; for instance, the RL and TM moved out of phase with the BM at low frequencies. The dependence of the phases on stimulus level and measurement location is described in detail later.

Average displacement gains and phases from a number of mice both alive and postmortem are shown in **Figure 2**, with the differences between the live and postmortem measurements shown in **Figure 3**. Death strongly reduces the endocochlear potential and any mechanotransduction currents, and thus diminishes OHC activity^31^. Consistent with this, displacement gains near the CF were greatly reduced postmortem and nonlinearity was nearly eliminated. However, while BM displacement gains an octave or more below the CF were little changed after death, the responses of the OHC-DC junction and RL were reduced at all frequencies, even for high stimulus levels. TM and LC region gains were also generally reduced postmortem, except for 90 dB SPL stimuli presented at ∼3 to 5 kHz. For a 1 kHz, 90 dB SPL stimulus, the average live re dead gain was 12 dB for the TM and ∼23 dB for the OHC-DC junction, RL, and LC region. The average data therefore confirm that all locations beyond the BM were influenced by active, broadband amplification.

**Figure 2.**
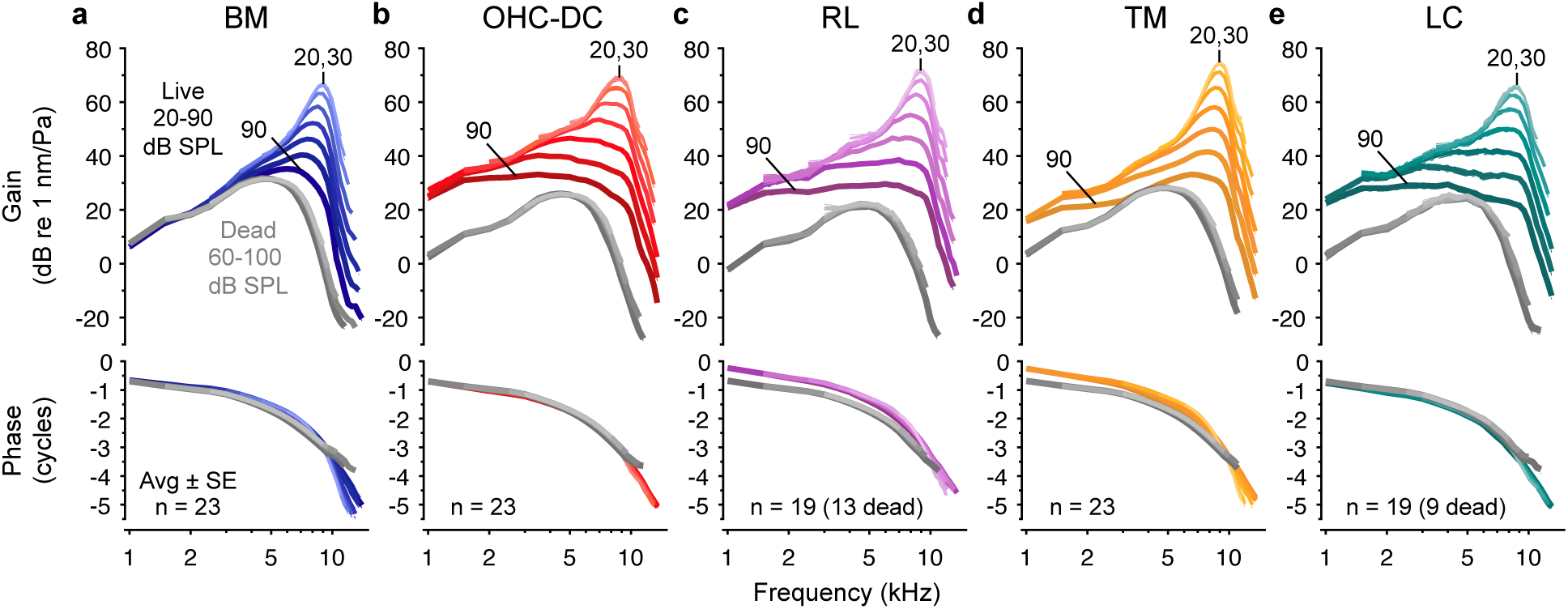
Low-frequency responses of all locations beyond the BM are actively amplified. (**a**-**e**) Average displacement gains (top panels) and phases (bottom panels) for all measurement locations in both live and dead mice. Responses are shown for levels of 20 to 90 dB SPL in live mice and 60 to 100 dB SPL in dead mice, with thicker and darker lines used for higher stimulus levels. For live mice, average gain curves for 20 and 30 dB SPL stimuli were similar and are indicated numerically, along with the average gain curve for 90 dB SPL stimuli. Postmortem responses largely overlapped, indicating their linear growth. BM, OHC-DC junction, and TM responses were obtained from all mice, while RL and LC region responses were obtained from smaller subsets. The low reflectivity of the RL and LC region particularly limited the ability to obtain postmortem measurements. Dashed-dotted lines indicate ±1 SE, though these are typically obscured by the average curves.

**Figure 3.**
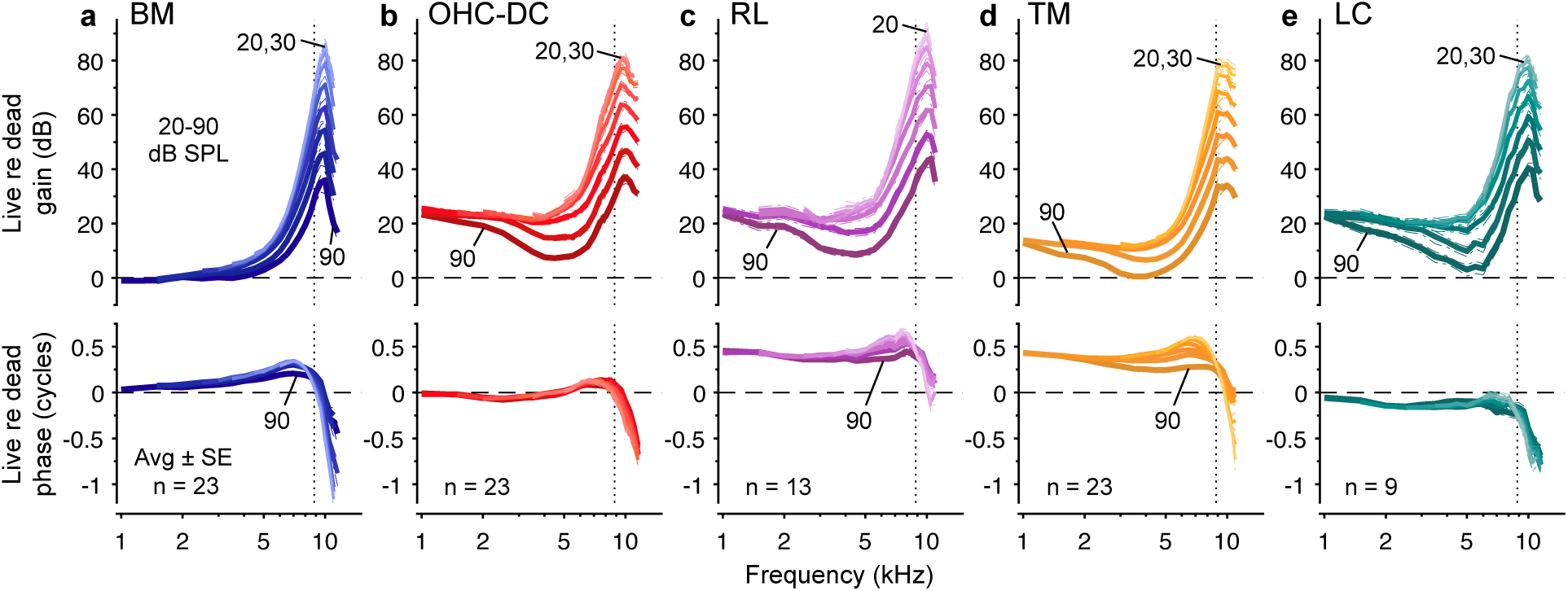
Gain and phase of live relative to postmortem responses. (**a**-**e**) Average displacement gains and phases in live mice relative to the postmortem response for each measurement location. Since displacements were essentially linear postmortem, responses for stimulus levels of 20 to 90 dB SPL in live mice were referenced to the postmortem response obtained at 100 dB SPL. Gain and phase differences at higher stimulus levels are shown with thicker and darker lines, with gain differences for the lowest and highest stimulus levels indicated numerically. Gain differences for 20 and 30 dB SPL stimuli were indistinguishable except for the RL. Phase differences were similar across stimulus level, though the phase leads observed for the BM, RL, and TM decreased slightly at higher stimulus levels for frequencies approaching the CF (the average CF of 8.8 kHz is indicated by vertical dotted lines). Dashed-dotted lines indicate ±1 SE.

Displacement phases also changed postmortem (bottom panels of **Fig. 2** and **3**), most notably for the RL and TM. At frequencies up to the CF, RL and TM displacement phases in live mice led those obtained postmortem by ∼0.5 cycles, with this phase lead being slightly smaller for higher stimulus levels and near-CF stimuli. BM displacement phases in live mice were similar to those obtained postmortem at low frequencies, while they exhibited a ∼0.25 cycle lead near the CF. OHC-DC junction displacement phases changed little after death despite this region’s motion being strongly influenced by electromotility in live mice. LC region displacement phases in live mice slightly lagged those obtained postmortem at most frequencies below the CF. Above the CF, displacements of all locations accumulated phase more rapidly with frequency in live mice than in dead mice.

### Relative motions are consistent with the influence of broadband OHC electromotility

In live mice, the relative displacement magnitudes and phases were consistent with the influence of cycle-by-cycle OHC length changes that were shaped by low-pass filtering^24^ (**Fig. 4a-e**). When normalized to BM motion, OHC-DC junction displacements declined by ∼6 dB/octave for low stimulus levels and frequencies below the CF (**Fig. 4a**, top panel). The OHC-DC junction also went from moving in phase with the BM at low frequencies to lagging the BM by ∼0.25 cycles at the CF (**Fig. 4a**, bottom). The RL followed a similar pattern except that it moved ∼0.4 cycles out of phase with the BM at low frequencies, with this phase lead decreasing to ∼0.2 cycles or less near the CF (**Fig. 4b**, bottom). The RL and OHC-DC junction therefore moved out of phase, consistent with OHC contraction and elongation at the stimulus frequency. Additionally, both the amplitude and phase behaviors appeared to reflect the influence of a first-order low-pass filter^24,32^, which may result from filtering of the receptor potential by the OHC membrane’s electrical properties^33^.

**Figure 4.**
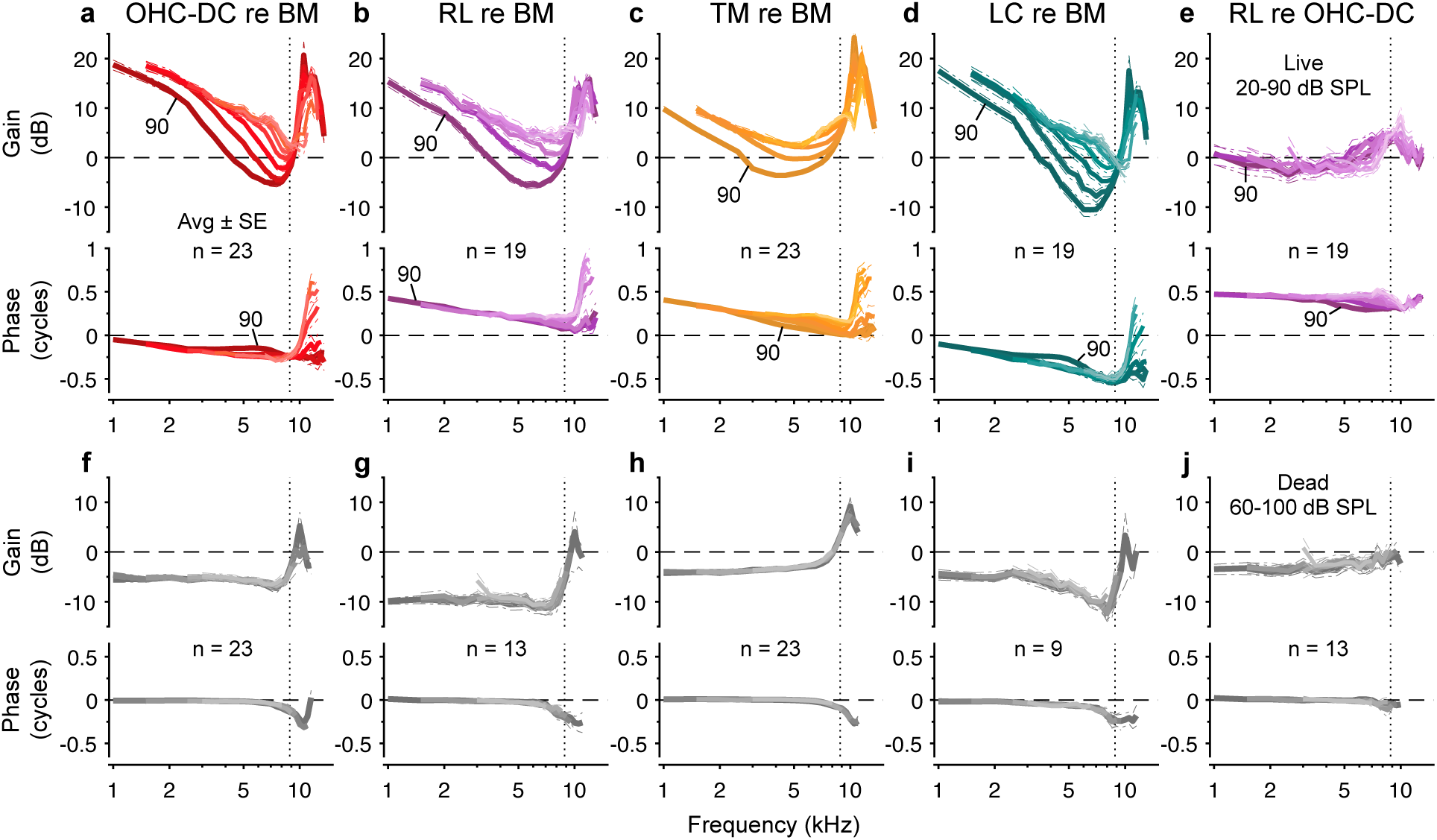
Relative motions are consistent with cycle-by-cycle OHC electromotility. (**a**-**d**) Average displacement gains and phases of the OHC-DC junction, RL, TM, and LC region relative to the BM in live mice for stimulus levels of 20 to 90 dB SPL. Below the average CF (indicated by vertical dotted lines), the relative gains and phases rolled off with frequency in a manner consistent with the influence low-pass filtering. (**e**) Average displacement gain and phase of the RL relative to the OHC-DC junction. The RL moved roughly out of phase with the OHC-DC junction. In **a**-**e**, relative responses at higher stimulus levels are shown with thicker and darker lines, with responses for 90 dB SPL stimuli also indicated numerically. (**f**-**j**) As in **a**-**e**, but showing the average relative responses in dead mice for stimulus levels of 60 to 100 dB SPL. In all panels, dashed-dotted lines indicate ±1 SE.

While motion of the TM relative to the BM exhibited less of a low-pass characteristic, the TM also led the BM by ∼0.4 cycles at low frequencies, with the phase lead decreasing to ∼0.15 cycles at the CF for low stimulus levels and ∼0 cycles for high levels (**Fig. 4c**). The TM presumably moves similarly to the RL because it is connected to the tallest row of OHC stereocilia. In contrast, the motion of the LC region relative to the BM revealed a stronger low-pass characteristic than the OHC-DC junction or RL, with a ∼0.5 cycle lag near the CF, as observed previously^24,34^ (**Fig. 4d**). The LC region generally moved out of phase with the adjacent RL, consistent with this region pivoting in the opposite direction when the OHCs exert force on the RL^35,36^.

The phase difference between OHC-DC junction and RL motion was closest to 0.5 cycles at low stimulus levels and low frequencies (**Fig. 4e**). At high stimulus levels and frequencies near the CF, the phase difference likely diminishes due to the increasing influence of the underlying traveling-wave motion. This is indicated by the motion phases becoming more similar to the BM phase at high stimulus levels, and the motion magnitudes becoming similar to or smaller than BM motion. Above the CF, the influence of traveling-wave motion appears to decrease, such that all locations move much more than the BM, regardless of stimulus level.

Postmortem, the BM moved more than the other locations though the motion phases were similar for all locations below ∼7 kHz (**Fig. 4f-i**). At higher frequencies, motions at locations beyond the BM lagged the BM by ∼0.25 cycles and became more similar in magnitude to BM motion, if not larger. Importantly, the phase difference between RL and OHC-DC junction motion was eliminated postmortem, and the RL moved slightly less than the OHC-DC junction, except near the CF (**Fig. 4j**).

### Compressive and non-monotonic growth at low frequencies is revealed at high stimulus levels

As little compression was observed at the lowest frequencies tested for stimulus levels up to 90 dB SPL, higher levels were used to determine whether any nonlinearity could be elicited at these frequencies. Indeed, data for an individual mouse in **Figure 5** show that increasing the stimulus level from 90 to 100 dB SPL reduced the gain of OHC-DC junction and TM responses even below 2 kHz, bringing them closer to those obtained postmortem. BM motion, in contrast, remained largely linear.

**Figure 5.**
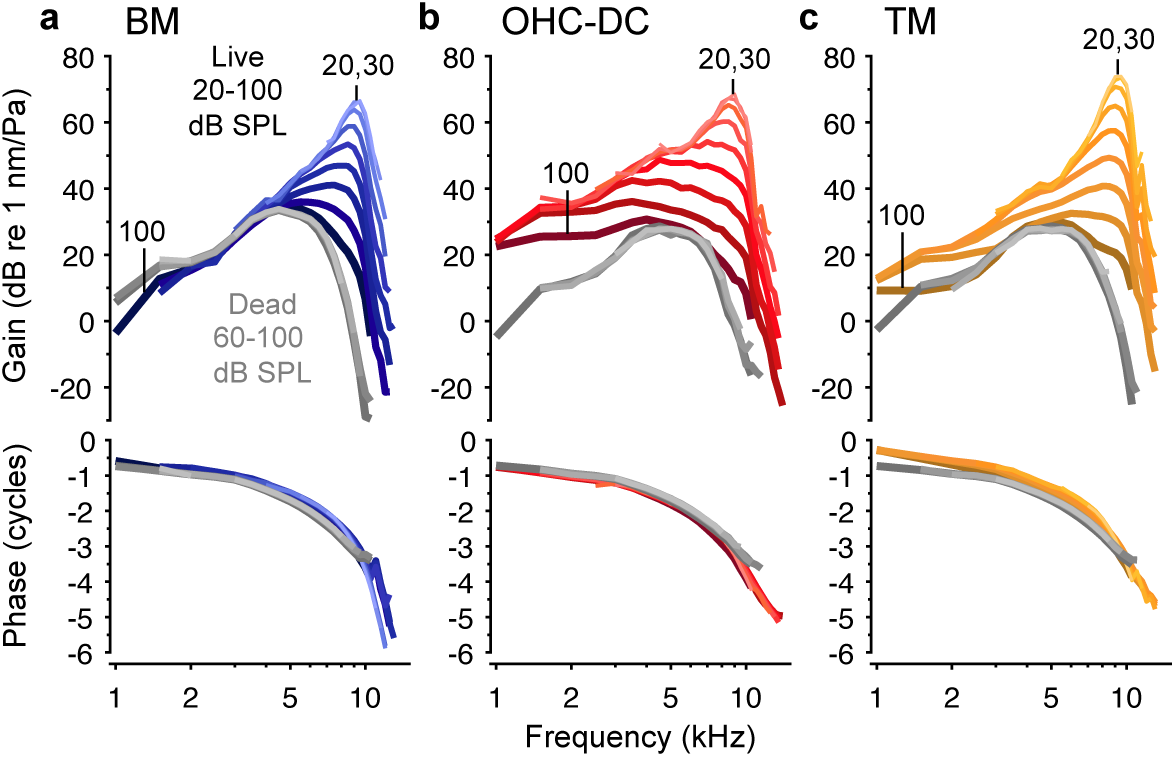
Low-frequency extent of nonlinearity broadens with increasing stimulus level. (**a**-**c**) Displacement gains and phases for the BM, OHC-DC junction, and TM at stimulus levels of 20 to 100 dB SPL in an individual mouse, both alive and postmortem. At low frequencies, increasing the stimulus level from 90 to 100 dB SPL in the live condition caused little change to BM responses but reduced the OHC-DC junction and TM displacement gains. The phases also developed an additional lag at high frequencies for 100 dB SPL stimuli, though these changes are obscured by the postmortem phases. Responses for increasing stimulus levels are shown with thicker and darker lines, and gains for the lowest and highest stimulus levels are indicated numerically.

To better characterize the nonlinear growth at low frequencies, responses to 1, 1.5, and 2 kHz tones were examined at levels of up to 115 or 120 dB SPL. As illustrated by data from an individual mouse (**Fig. 6a-c**), BM responses grew near-linearly at all levels, while the motions of the other locations exhibited compressive growth at levels that depended on both stimulus frequency and location. OHC-DC junction and LC region responses grew smoothly with level and were qualitatively similar, with the most compressive growth occurring at 90 to 100 dB SPL and milder compression at higher levels. RL and TM responses exhibited the strongest compression, at least for levels below ∼100 dB SPL, with magnitude notches followed by linear or expansive growth at higher levels. Magnitude notches were consistently observed for the TM while their presence in RL responses was variable, as demonstrated by RL responses from two other mice (**Fig. 6d**). When present, notches in RL responses also typically occurred at higher stimulus levels. Average magnitude slopes for each location are compared for 1.5 kHz stimuli in **Figure 6e**, confirming the trends described above.

**Figure 6.**
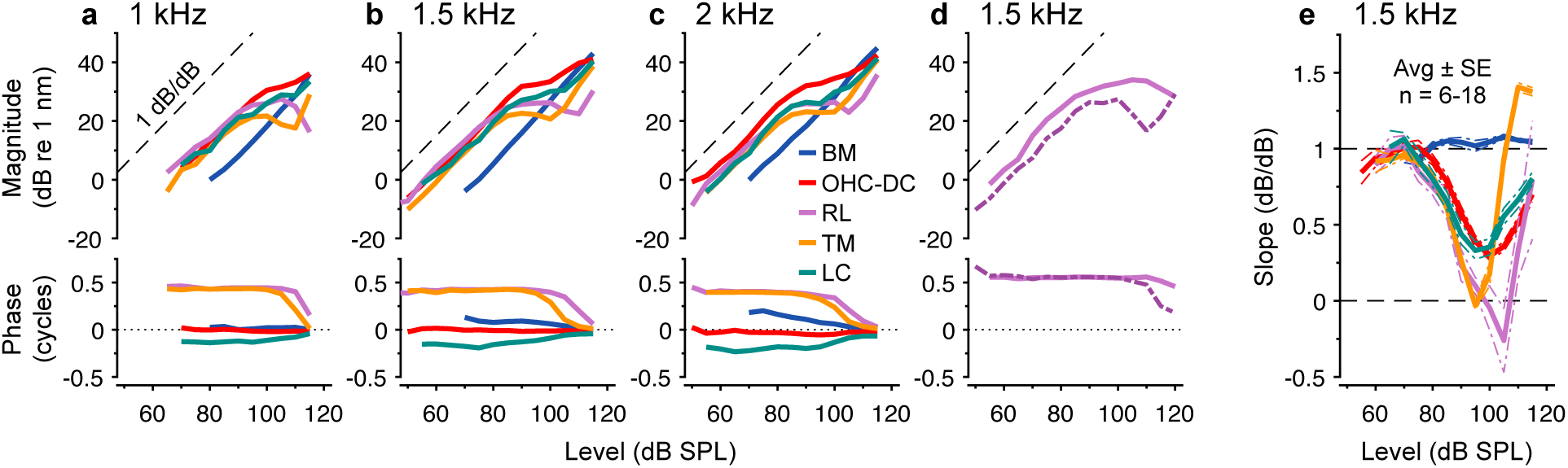
Low-frequency responses exhibit variable growth patterns and converge in phase at high stimulus levels. (**a**-**c**) Displacement magnitudes and phases for all locations in response to 1, 1.5, and 2 kHz tones varied in level from 0 to 115 dB SPL in 5 dB steps for an individual mouse. RL and TM responses grew most compressively and often exhibited decreasing magnitudes or notches at high stimulus levels. Phases were referenced to the BM response phase at 115 dB SPL and became similar at the highest levels. (**d**) RL responses to 1.5 kHz tones presented at levels of up to 120 dB SPL for two other mice, illustrating variability in the occurrence of notches and associated phase shifts. (**e**) Average growth slopes for responses to 1.5 kHz tones as a function of stimulus level from 18 mice (for the BM, OHC-DC junction, and TM) or different subsets of 9 mice (for the RL and LC region). Responses were obtained at levels of up to 115 and/or 120 dB SPL in 5 dB steps, with slopes averaged across both measurements for each mouse, when available. Slopes were obtained using the linear-regression fit to the three-point span centered at each stimulus level and were only calculated for responses that were above the measurement noise. Average slopes are shown for a given stimulus level if data were available from at least six mice. Dashed-dotted lines indicate ±1 SE.

Though the RL and TM moved roughly out of phase with the OHC-DC junction at most stimulus levels, the phases of all locations tended to converge at the highest levels tested (see bottom panels in **Fig. 6a-c**). Shifts in RL and TM motion phases occurred at roughly the levels where the magnitude notches were observed, suggesting that these features were due to interference between the active OHC-induced motion and the underlying traveling-wave motion, with the latter becoming dominant at high levels. In accordance with this, when no notches were observed in RL motion, the phases were more constant across level, indicating that the active motion remained dominant (see data shown with solid lines in **Fig. 6d**). While the OHC-DC junction motion phase varied little with level and was similar to the high-level “passive” motion, BM motion sometimes exhibited frequency- and level-dependent phase leads, as in the example shown. In other mice, the BM moved with the OHC-DC junction or exhibited a constant phase lead. This variability could be due to differences in the radial position of the measurement along the BM and may belie some influence of OHC electromotility at low stimulus levels, despite the approximately linear growth.

To illustrate the relation between the active and truly passive motions, average displacement gains and phases for 1.5 kHz tones presented at levels up to 120 dB SPL are shown for live and dead mice in **Figure 7**. In live mice, all locations beyond the BM exhibited nonlinear gain at levels above 80 dB SPL. At higher levels, the gains and phases became more similar to those measured postmortem. For the TM, the live gain actually decreased below the postmortem gain at levels where a notch in the live response occurred. This confirms that these notches result when the out-of-phase active and passive motions become similar in magnitude, such that they destructively interfere. Notches and associated phase shifts were not as obvious in the average RL responses due to the variability in these features, and the average RL motion phase in live mice did not fully converge with the average postmortem phase. This is consistent with the RL’s active motion tending to dominate over a wider range of stimulus levels, when compared to the TM. The RL’s maximum average live gain was almost 10 dB greater than that of the TM, while its passive or postmortem gain was ∼6 dB lower.

**Fig. 7.**
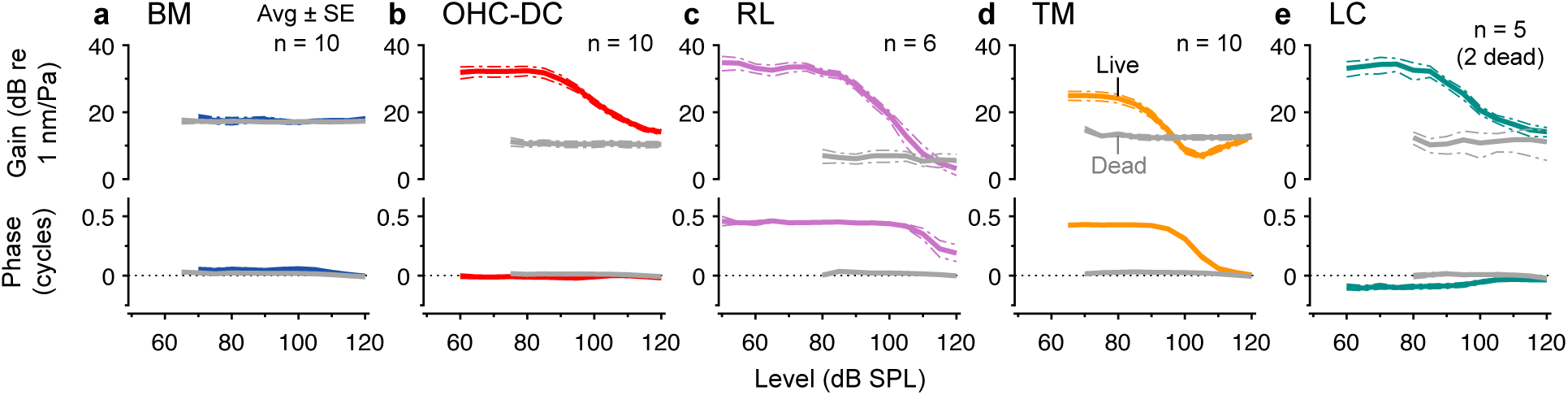
At high stimulus levels, low-frequency responses in live mice become similar to the passive traveling-wave motion. (**a**-**e**) Average displacement gains and phases for all locations in response to 1.5 kHz tones varied from 0 to 120 dB SPL in live and dead mice, with the latter shown in gray. All phases were referenced to the BM response phase in live mice at 120 dB SPL. Live and postmortem responses were obtained from the BM, OHC-DC, and TM in the same 10 mice, with different subsets contributing to the average RL and LC region responses. Postmortem RL responses are from four of the six mice in which live responses were obtained, as well as two other mice. Postmortem LC region responses are from two of the five mice in which live responses were obtained. Dashed-dotted lines indicate ±1 SE.

### Low-frequency responses can be suppressed by a second tone

In addition to growing compressively at high stimulus levels, low-frequency responses could also be suppressed by a second tone at a different frequency. This mechanical two-tone suppression is thought to result from the second tone saturating the mechanotransduction currents, so that less current is generated in response to the first tone^37^. As shown by data from an individual mouse in **Fig. 8a**, TM responses to an 80 dB SPL, 1.5 kHz tone could be strongly and non-monotonically influenced by presenting a 7 kHz tone that was varied in level. As the suppressor level increased above 50 dB SPL, the 1.5 kHz response rapidly dropped but then increased again before becoming relatively stable for suppressor levels of 80 dB SPL and higher. This non-monotonic behavior resembles the notches observed in TM responses measured as a function of stimulus level, which were likewise associated with a ∼0.4 cycle phase shift (e.g., **Figs. 6** and **7**). Increasing the suppressor tone level to ∼70 dB SPL therefore appeared to reduce the active motion at 1.5 kHz to where it was similar in magnitude to the passive motion, resulting in maximal cancellation. Further reduction of the active response at higher suppressor levels then revealed the passive motion.

**Figure 8.**
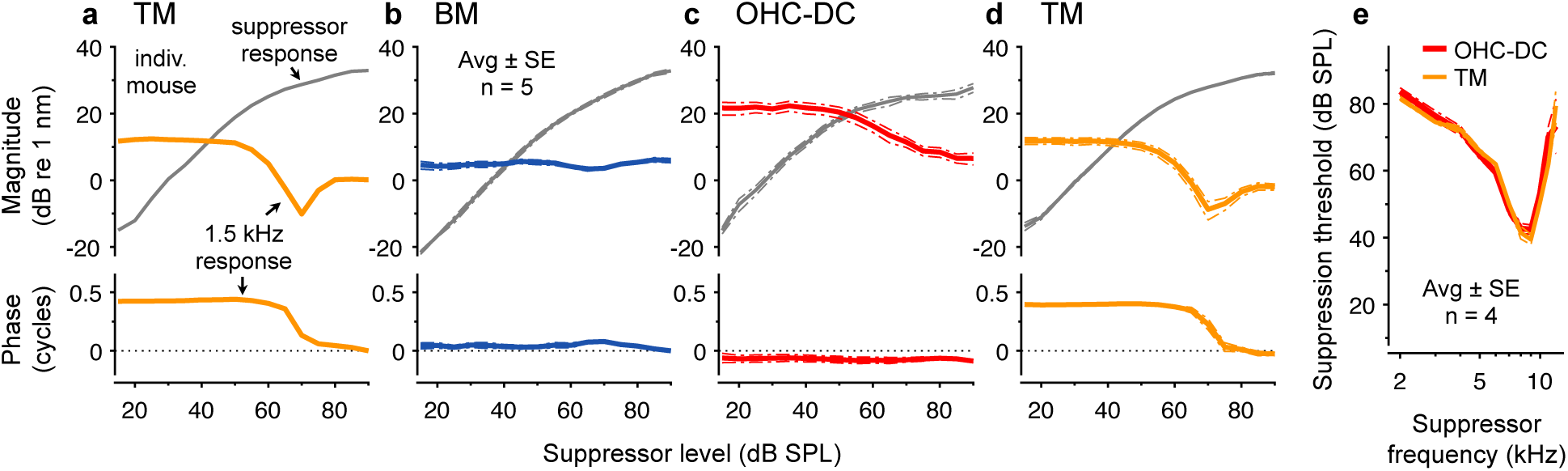
Low-frequency responses of the TM and OHC-DC junction are suppressed by a second tone. (**a**) Magnitudes and phases of TM displacements in an individual mouse measured in response to a 1.5 kHz tone presented at 80 dB SPL along with a 7 kHz suppressor tone that was varied in level. Increasing the level of the 7 kHz tone produced non-monotonic changes in the magnitude of the 1.5 kHz response as well as a ∼0.4 cycle phase shift. The magnitude of the response to the 7 kHz tone is shown in gray. (**b**-**d**) Average magnitudes and phases of BM, OHC-DC junction, and TM displacements measured in five mice using the same stimulus paradigm as in **a**. In contrast to the complex amplitude and phase changes observed in TM responses to the 1.5 kHz tone, BM responses were only slightly altered while OHC-DC junction responses declined smoothly as the suppressor level increased. In **a**-**d**, phases were referenced to the BM response phase when the suppressor was 90 dB SPL. (**e**) Average level of the suppressor tone required to reduce OHC-DC junction and TM responses to an 80 dB SPL, 1.5 kHz tone by a threshold criterion of 1.5 dB for suppressor frequencies ranging from 2 to 12 kHz. In **b**-**e**, dashed-dotted lines indicate ±1 SE.

Non-monotonic suppression was observed across all mice for TM responses, but not for responses of the BM or OHC-DC junction, as shown by average data in **Fig. 8b-d**. As expected, BM responses at 1.5 kHz were minimally affected by a 7 kHz tone, though small amplitude and phase variations could be observed for suppressor levels of ∼60 to 70 dB SPL. OHC-DC junction responses declined smoothly as the 7 kHz tone was increased in level but underwent little change in phase, consistent with the active and passive OHC-DC junction motions having similar phases (see **Fig. 7b**). While increasing the suppressor level caused more rapid reduction of the 1.5 kHz response for the TM compared to the OHC-DC junction, the response of both locations could be reduced by a fixed, small amount using roughly the same suppressor level. The average suppressor level required to reduce OHC-DC junction and TM responses by 1.5 dB in four mice is plotted as a function of suppressor frequency in **Fig. 8e**. The suppression threshold curves are similarly tuned and largely overlap, indicating that the same mechanisms are involved in suppression of low-frequency motion at the TM and the bottom of the OHC region.

### Compression and suppression of low-frequency responses can be replicated with a simple model

A simple model was used to approximate the behavior of low-frequency responses of the OHC-DC junction and the TM, with the total motion of each location considered to be the vector sum of active and passive motion components (**Fig. 9**). A similar approach has previously been used to replicate the level-dependent growth of displacements at the stimulus frequency and any associated distortions^24,38,39^.

**Figure 9.**
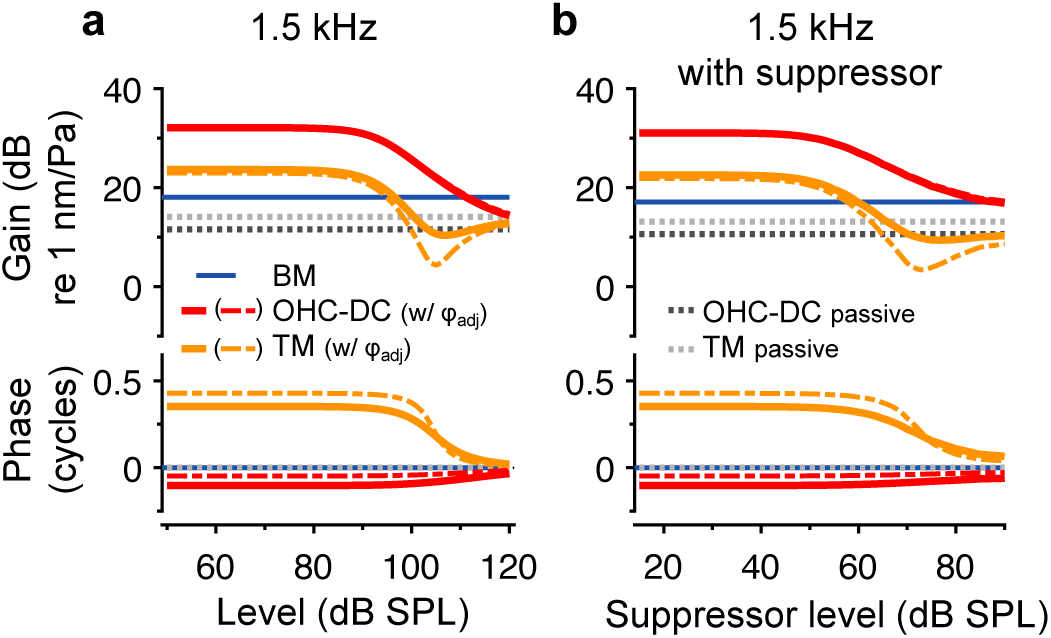
Growth and suppression of low-frequency responses can be replicated with a simple model. (**a**) Modeled displacement gains and phases for BM, OHC-DC junction, and TM responses to a 1.5 kHz tone varied in level. BM motion was assumed to be passive and linear, while OHC-DC junction and TM responses were approximated by summing the estimated active and passive motion components. The passive components (shown with dotted lines) were scaled versions of BM motion. The active components were proportional to the low-pass filtered output of a Boltzmann function with BM displacement used as the function’s input. The phase of the TM’s active motion component was then rotated by 0.5 cycles. The model output after adjusting the active component phase by adding 0.06 cycles is also shown with dashed-dotted lines (labeled “w/ φ_adj_” in the legend), demonstrating the influence of phase on the depth of the notch in TM gain. All phases are referenced to the BM motion phase. (**b**) As in **a**, but showing the modeled displacement gains and phases for a 1.5 kHz, 80 dB SPL tone presented with a 7 kHz suppressor tone that was varied in level. For clarity, modeled responses to the 7 kHz suppressor are not shown.

In the model, the active motion was proportional to the low-pass filtered output of a first-order Boltzmann function. This function was used to represent the nonlinear relation between stereociliary bundle deflection (*x*) and mechanotransduction current (*I*) and is defined as

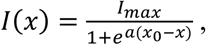

where *I*_max_ is the maximum current (here normalized to 1), *a* is a slope parameter (here 0.28 nm^-1^), and *x*_0_ is the bundle position where the current is half activated (here 2.6 nm). For simplicity, BM displacements were used as a proxy for stereociliary displacement, with the average BM response to the highest stimulus level linearly scaled to estimate the input for each desired stimulus level. The Boltzmann’s output was passed through a first-order low-pass filter with 1.75 kHz corner frequency to approximate the influence of OHC membrane filtering on the receptor potential. The resulting quantity was then scaled to replicate the active motion induced by OHC motility at the OHC-DC junction and TM (by factors of 100 and 50, respectively). The active TM motion was further assumed to lead the OHC-DC junction motion by 0.5 cycles. The passive motions of the OHC-DC junction and TM were estimated by scaling BM displacement by factors of 0.47 and 0.63, respectively, which are roughly the ratios of OHC-DC junction and TM motion to BM motion observed postmortem.

The gains and phases of the modeled OHC-DC junction and TM responses to 1.5 kHz tones varied in level (**Fig. 9a**) resembled those shown in **Figure 7**, particularly after adding 0.06 cycles to the phases of the active component (see dashed-dotted lines). This phase adjustment, which partly counteracted the effects of low-pass filtering, produced response phases that better matched the data and enhanced the phase difference between the active and passive TM motion components, resulting in stronger interference at high stimulus levels. The model could also approximate responses to an 80 dB SPL, 1.5 kHz tone measured in the presence of a 7 kHz suppressor tone (compare **Fig. 9b** to **Fig. 8**). This confirms that compression and suppression of low-frequency responses can be attributed to the same saturating nonlinearity and that the non-monotonic behavior of TM responses arises from interference between active and passive motion components.

## DISCUSSION

The present report clearly demonstrates that the motions near the top of the organ of Corti, including at the RL and TM, are influenced by nonlinear, broadband OHC force generation in the apex of the mouse cochlea. This was revealed by the physiological vulnerability and nonlinear growth of motions at frequencies far below the CF, as well as the suppressibility of low-frequency TM motions by a second tone. Importantly, the below-CF nonlinearity observed in RL motion could not simply be due to contamination by the motion of the OHC-DC junction, as the movements were in opposite directions. Low-frequency nonlinearity was also observed for measurement points on the TM that were too distant from the OHC-DC junction to be influenced by such artifact. The present measurements therefore confirm prior observations of low-frequency nonlinearity in mouse cochlear apex^18,20,24,28,29,39^ with greater spatial certainty.

Evidence for broadband, nonlinear amplification of RL motion has also been previously reported for more basal locations by Ren and colleagues, with the motions of the ∼45 kHz region of the mouse^19^ and ∼20-25 kHz region of the gerbil^22^ being observed through the round window membrane. In gerbil, broadband nonlinearity was also present at the RL when the same CF location was viewed through a cochleostomy, which allowed measurement of the primarily transverse motion^40^. Regardless of viewing angle, the RL moved roughly out of phase with the BM at low frequencies. Thus, while their approach also did not allow the precise identification of the RL’s axial location, the reported low-frequency nonlinearity and phase behaviors are consistent with those observed here in the mouse apex.

Nevertheless, these measurements contrast with several other reports. Using an OCT system capable of resolving individual OHCs when viewed through the round window, Cho and Puria^26^ found little nonlinearity or physiological vulnerability in low-frequency motions of the RL measured from the 45 kHz region in gerbil. Similar findings in gerbil were reported for locations with CFs of 25 to 45 kHz by Strimbu et al.^27,41^, who used a lower-resolution system to observe motions through either the round window or a cochleostomy. Measurements by the same group from the base of the guinea pig cochlea revealed mild low-frequency nonlinearity in RL motion when viewed through the round window, but variable results when viewed through a cochleostomy^42,43^. Interestingly, even when low-frequency RL motion is found to be linear, it often leads BM motion by up to 0.5 cycles, with this phase difference disappearing postmortem^26,43^. If this phase behavior reflects the influence of OHC electromotility, it is unclear how RL motion would remain un-amplified.

Measurements from apical locations in chinchilla, gerbil, and guinea pig suggest that nonlinearity in the motions at both the top of the organ of Corti and the BM becomes more broadband as the CF decreases below a few kHz^44–47^. It remains uncertain whether this a unique feature of low-CF regions or if the frequency extent and spatial distribution of the nonlinearity smoothly expand from the base to apex. Further work is needed to reconcile these diverse findings and understand how below-CF nonlinearity may depend on species, cochlear location, and viewing angle.

Though low-frequency OHC force generation influences the motion at all locations beyond the BM in the mouse apex, the degree of nonlinearity and the overall frequency response still varied across locations. For instance, from the TM to the OHC-DC junction, the responses became progressively more low-pass in character (e.g., **Fig. 4**). Additionally, LC region motion seemed slightly more low-pass than OHC-DC junction motion, with a lag re BM motion of ∼0.5 cycle at the CF, compared to the 0.25 cycle lag observed at the OHC-DC junction. These variable features could reflect differences in tissue properties or the relative influence of the active and passive motion components that align with the optical measurement axis.

The spatial variations described above are consistent with measurements at higher-frequency locations in both gerbil and guinea pig^26,27,43^. However, OHC-DC junction motion at these basal sites often appears even more low-pass, particularly at high stimulus levels and in measurements that mostly capture the transverse motion. In some cases, this may be partly due to the active OHC-DC junction motion lagging BM motion by up to 0.5 cycles near the CF^26,27^. The traveling-wave motion transmitted by the BM can therefore interfere with the active motion so that the total displacement declines at frequencies approaching the CF. While LC region motion in the mouse apex also lagged BM motion by ∼0.5 cycles near the CF, interference may not entirely explain the low-pass character of these responses in live mice, as even the postmortem motion of the LC region relative to the BM declined above ∼2 kHz.

By amplifying RL and TM motion at frequencies far below the CF, OHCs may have a direct, local influence on IHC stimulation at these frequencies^13,30^. This below-CF amplification could account for phenomena that have been observed in auditory nerve responses in other species. For instance, for high-CF auditory nerve fibers in cats, responses to below-CF stimuli can be reduced by tones^30^ and electrical stimulation of the medial olivocochlear efferent fibers, which inhibit OHC activity^48,49^. Such inhibition could conceivably be due to an efferent-induced reduction in the RL and/or TM motions that are involved in stimulating the IHCs. Even though low-frequency responses of high-CF IHCs are small compared to those of the OHCs^50^, these responses could still be driven by amplified motion that can be inhibited or suppressed. The IHC response to amplified low-frequency motion may be attenuated by a variety of mechanisms – e.g., sensitivity to velocity rather than displacement^13,51^.

The notches and phase shifts observed in motions of the RL and TM could also underlie notches and associated phase shifts in auditory nerve fiber responses that occur above ∼90 dB SPL^52–55^. Such notches have primarily been examined at the low frequencies where nerve fibers phase-lock to the stimuli and appear to have correlates in IHC receptor potentials^56^. They have long been proposed to result from interference between two modes of IHC stimulation: an active, physiologically vulnerable mode at low levels and a more passive mode at high levels^53,54^. The present data suggest that these modes may simply correspond to the active and passive motions of the RL and/or TM. However, it remains to be determined if OHCs provide broadband amplification of RL and TM motion in the species for which notches in nerve fiber responses have been observed. Additionally, whether these notches are related to RL or TM motion depends on how these motions interact to stimulate the IHCs, which is incompletely understood^13,57^.

Given that the tallest row of OHC stereocilia extend from the RL to the TM, it is also plausible that below-CF amplification and related interference effects could influence the OHC’s own stimulation. While notches and phase shifts have been observed in OHC receptor potentials measured at high-CF regions in guinea pig, they were primarily found within a half-octave above or below the CF and exhibited a complex frequency-dependence^58,59^. Such features therefore seem unlikely to result from interference between the OHC’s active and passive motions, since the relative phases of these motions should vary more gradually with frequency. Further technological advances are needed to conduct the *in vivo* measurements of stereocilia motion that are required to fully understand the relationships among these various phenomena.

## METHODS

### Mice

Measurements were obtained from 3- to 7-week-old CBA/CaJ mice bred and housed at the University of Southern California. A total of 23 mice (11 female) were used. All procedures were approved by the local Institutional Animal Care and Use Committee and have been detailed previously^24^.

Briefly, mice were deeply anesthetized (80-100 mg/kg ketamine and 5-10 mg/kg xylazine) and placed on a heating pad set at 38 °C. Supplemental anesthesia was administered to maintain areflexia throughout the experiment. After fixing the skull to a head holder with dental cement, the left bulla was surgically accessed and carefully opened to provide a view of the cochlear surface and ossicular chain. The pinna and a portion of the ear canal were removed so that an acoustic probe containing speakers and a microphone (Etymotic Research ER-10X; Elk Grove Village, IL) could be sealed over the eardrum with dental cement. A tracheotomy was also performed to facilitate free breathing. After all desired *in vivo* measurements were completed, mice were euthanized via anesthetic overdose and certain measurements were repeated postmortem.

### OCT imaging and vibrometry

Vibrations of the cochlear structures and middle ear ossicles were measured with a custom-built, swept-source OCT system that has been described previously^24^. The system uses a light source with 1310 nm center wavelength and 95 nm bandwidth (Insight; Broomfield, CO) and provides axial and lateral imaging resolutions of 11 and 9.8 µm (full width at half maximum), respectively. The actual axial and lateral dimensions of the pixels in the images that were selected as measurement points were ∼4.8 µm.

After scanning the light source across the cochlear bone to obtain two-dimensional, cross-sectional images of the apical turn, sound-evoked displacements were measured from the BM, OHC-DC junction, RL, TM, and LC region. The mouse’s head was positioned to minimize the angle between the BM and the horizontal plane, which fell between 10 and 30° and was ∼20° on average. The measured displacements therefore primarily captured the transverse motions of the structures, though they likely also included some mixture of radial and longitudinal motions. Indeed, OHCs are tilted in both the radial and longitudinal directions^60^. The CF of the measurement site was defined by the frequency of the peak BM response for 30 dB SPL stimuli and fell between 8.5 and 9.5 kHz.

Measurements were always made from points that were associated with local peaks in reflectivity when examined as a function of axial distance. This maximized the signal-to-noise ratio and reduced potential contamination by signals reflected from adjacent locations^61^. As the reflectivity could fluctuate over time, cross-sectional images were obtained between vibration measurements to assess which points might yield high-quality data. The reflectivity of the RL and LC region was often low and/or unstable, thus limiting success in recording from these locations and reducing the number of mice contributing data to the average responses.

### Stimulus paradigms

Acoustic stimuli were calibrated using the pressure measured by the probe microphone after correcting for its frequency-dependent sensitivity. Displacement responses were obtained from each measurement point using discrete tones swept in frequency from 1 to 15 kHz in 0.5 kHz steps and varied in level from 10 to 90 dB SPL in 10 dB steps. In a few mice, frequency responses were also obtained at levels up to 100 dB SPL just prior to euthanasia. Stimuli were 102 ms long (including 1 ms cosine-squared ramps) and presented once every 110 ms. Responses were typically averaged over 4 or 8 stimulus repetitions. For points with unstable reflectivity, responses were sometimes obtained for a single stimulus presentation to minimize changes in reflectivity during the measurement.

The level-dependence of displacements at specific low frequencies was also examined from 0 to 115 or 120 dB SPL in 5 dB steps, with responses averaged over 4 stimulus repetitions. Measurements eventually focused on characterizing the growth of responses at 1.5 kHz for levels up to 120 dB SPL. *In vivo* displacement measurements from the ossicular chain for each stimulus paradigm were obtained in at least three mice to verify the absence of nonlinearity or phase shifts in the mechanical input to the cochlea. BM responses to 30 dB SPL tones presented at the CF also showed no consistent changes in sensitivity after the measurements, suggesting that the high sound pressures used did not cause any acoustic trauma. This is likely because the mouse middle ear and cochlea are not designed to efficiently transmit low-frequency stimuli^62^. Higher-frequency stimuli were not presented at levels exceeding 100 dB SPL due to the potential for such trauma as well as limitations of the acoustic probe’s output.

Two-tone stimuli were used to study how the response to a 1.5 kHz tone was affected by a second, “suppressor” tone. The 1.5 kHz tone was presented for 104 ms with a suppressor tone of equivalent duration turned on after 52 ms. Both tones had 2 ms onset and offset ramps, resulting in three 50 ms intervals containing the steady-state response to the 1.5 kHz tone alone, both the 1.5 kHz tone and suppressor tone, and the suppressor tone alone. The 1.5 kHz tone was presented at 80 dB SPL while the suppressor tone varied from 15 to 90 dB SPL in 5 dB steps. Suppressor frequencies were typically swept from 1 to 15 kHz in 1 kHz steps, though a more limited range of frequencies was sometimes used.

For all stimulus paradigms, a fast-Fourier transform was applied to the steady-state portion of the final response waveform to obtain the magnitude and phase of the displacement at the relevant frequencies. To estimate the measurement noise, the mean and standard deviation of the displacements at frequencies within 40 to 140 Hz below or above the response frequency were computed. Responses with magnitudes less than the mean plus three standard deviations of the displacements in surrounding frequency bins were considered to be below the measurement noise. Such responses were not plotted or included in subsequent averages or analyses.

### Postmortem measurements

All postmortem measurements were obtained within 50 minutes after death, as responses can change at later times^29^. Due to the lower and less stable reflectivity of the RL and LC region, it was difficult to record from these locations for all stimulus paradigms within this time constraint. Postmortem data from these locations were therefore only available from a subset of mice.

### Data reporting

Reported displacement magnitudes are peak values and displacement phases were referenced to the stimulus phase measured in the ear canal. Plots of average data show the mean ± 1 standard error (SE). Averages only include responses meeting the signal-to-noise criterion. Unless otherwise noted, averages are only shown for a given stimulus frequency or level if responses met the signal-to-noise criterion in at least two-thirds of the mice tested. While frequency responses were obtained down to levels of 10 dB SPL, these data are omitted here for visual clarity. Responses to 10 dB SPL tones were typically only above the measurement noise for a narrow range of frequencies and had displacement gains similar to those of responses to 20 and 30 dB SPL tones^24^. To reduce visual clutter, isolated data points or segments that were just above the measurement noise were also omitted from some plots.

## ACKNOWLEDGMENTS

This work was supported by NIH/NIDCD R01 DC021006.

## Notes

### Competing Interest Statement

The authors have declared no competing interest.

